# The Batch-Resourcing Angiogenesis Tool (BRAT) to enable high-content microscopy screening of microvascular networks

**DOI:** 10.1101/2025.01.24.634836

**Authors:** Harriet Krek, Ashley R. Murphy, Ryan McKinnon, Rose Ann Franco, Mark C. Allenby

## Abstract

Vessel forming assays are a valuable *in vitro* technology to evaluate the vasculogenic and angiogenic potential of different cell types, matrix proteins, and soluble factors. Recent advances in high-content microscopy allow for vascular morphogenesis assays to be captured in real-time and in high-throughput screening purposes. Unfortunately, existing microvascular network quantification algorithms are either inaccurate, not user-friendly, or manually analyse one image at a time, unfavourable to high-content screening applications. This manuscript introduces BRAT, the Batch-Resourcing Angiogenesis Tool, a computer algorithm with an open-source graphic user interface to efficiently segment, skeletonize, and analyse large batches of vascular network images with high accuracy. Benchmarked across diverse clinical and cultured microvascular network images, BRAT is the most sensitive vascular network image analysis tool (94.5%) and exhibits leading accuracy (93.3%). BRAT’s multi-threaded processing automatically analysed 886 microscopy images at a speed of 0.17 seconds/image (2:29 minutes) on a performance computer or 2.31 seconds/image (34:04) on a typical laptop. This is 10-to-100 fold more time-efficient than existing tools, which require 12 to 16 seconds of direct user input per image. BRAT is broadly useful to analyse vessel networks of different endothelial cell types cultured on 2D substrates and within 3D biomaterials. BRAT represents a powerful approach for the accurate and high-content screening of vessel forming assays for disease models, regenerative medicines, and therapeutic testing.

## 1. Introduction

Biological and medical imaging is essential in evaluating cell and tissue structure during regeneration and disease [1]. Tissue function and dysfunction depends on nutrient supply provided by microcapillaries throughout the body, grown during morphogenic processes of vasculogenesis, angiogenesis, and arteriogenesis [2]. The growth, presence, or damage of microcapillaries are routinely imaged and analysed by clinical radiology or pathology departments [1]. Recently, vasculogenic cell culture assays are increasingly used to understand how microcapillary networks form, and how different patient cells, environments, or pharmaceutical therapies can support or impair microcapillary development or function [2,3]. Until recently, our ability to statistically compare microvascular network quantity or quality was limited to manually scoring biomedical images, creating bottlenecks in the high content screening of many drug targets and mechanistic studies of cardiovascular therapeutics [4].

The ability for computers to automatically and quantitatively analyse image features, computer vision [5], has transformed the way biological image analysis is performed, with applications across microscopy and biological discovery [6]. Digital pathology, an emerging field of research, promises faster and more accurate patient diagnosis [7], providing accurate identification and analysis of single-cells in 2D histology samples, for example, cancer cells to classify tumour regions [8]. But advances in digital pathology remain less translated for more complex tissue structures such as microcapillaries, necessary for microvascular network quantitation to be accurate, objective, and consistent. Cardiovascular disease remains the most common cause of death worldwide [9], and despite the rise of high content drug screening technologies to discover new treatments for diabetes [10], neurovascular diseases [11], and peripheral artery disease [12], there exists an unmet need to analyse microvascular network quality in large batches of microscopy images.

In 2005, AngioQuant was the first quantitative and standardised algorithm for microvascular network image analysis by segmenting interconnected pixels from microscopy images of angiogenic assays [13]. In 2011, AngioTool became a popular, light-weight, user-friendly software to analyse microvascular networks in microscopy images of cell cultures and tissue explants [14], and became recommended by the National Institute of Health (NIH) consensus guidelines as a standard software for angiogenesis assay interpretation [15]. Also in 2011, the Rapid Analysis of Vessel Elements (RAVE) was released with a graphical user interface [16]. The Rapid Editable Analysis of Vessel Elements Routine (REAVER) was published in 2020 with leading accuracy (88 - 95%) and processing speed (15 – 60s per image), benchmarked to AngioQuant, AngioTool, and RAVE by releasing its own large dataset of manually segmented microvascular network images to support benchmarking [17].

These four algorithms (AngioQuant, AngioTool, RAVE, REAVER) have become the most frequently cited open-source microvascular network image analyses per year in literature. Other open-source 2D fluorescence microscopy algorithms include VESGEN 2D (2009; [18]), AutoTube (2019; [19]), Vessel Tech (2020; [20]), Q-VAT (2023; [21]), while 3D confocal microscopy analyses include QIBER3D (2022; [22]), Vessel Metrics (2024; [23]), and µVES (2023; [24]). OCT image analysis has OCTAVA (2021; [25]) and our team developed medical image (MRA) analyses for intracranial vasculature (2021; [26]. In addition, VascuViz (2022; [27]) provides multi-modal imaging solution. In addition, there exist commercial software (IMARIS) able to perform microvascular network analysis toolkits [28]. Despite the dozens of microvascular image analysis tools, the 4 most frequently used and cited algorithms remain AngioQuant, AngioTool, RAVE, and REAVER due to: (1) their reasonable accuracy in analysing popular fluorescent microscopy images, (2) their easy accessibility in a user-friendly open-source computer GUI or software, and (3) REAVER’s recent benchmarking of these four tools on a broad dataset of fluorescent microscopy images.

We here present the Batch-Resourcing Angiogenesis Tool (BRAT) for high-content screening of imaged microvascular networks by 2D fluorescence microscopy. BRAT is presented in an open-source easy-to-use format. BRAT demonstrates best-in-class accuracy, speed, and consistency when compared on the same images as AngioQuant, AngioTool, RAVE, and REAVER. Importantly, these other microvascular network analysis tools require a user to spend minutes to self-select brightness and contrast parameters for individual images, before initiating the analysis taking several more minutes per image. A significant advantage of BRAT is its ability to automatically standardise and batch-process thousands of images in an hour (sub-seconds per image) using graphics processing unit (GPU) multi-threading. We demonstrate BRAT is the most accurate microvascular network analysis software to date and is uniquely capable to deliver high content screening analysis of vascular morphologies for live-imaging and drug screening.

## 2. Materials and Methods

### 2.1. Cell Culture and Microscopy

The use of primary umbilical cord blood and immortalised commercial cell lines was approved by UQ’s Human Research Ethics Committee in 2021/HE002698 and 2022/HE001310 and Institutional Biosafety Committee in IBC/602E/ChemEng/2023 and IBC/568B/ChemEng/2022.

Vasculogenic cell culture images were either prepared by our laboratory following the below protocols (**Fig. 1-3, 5**) or collected from the REAVER publication dataset for benchmarking (**Fig. 4**) [17]. All cultures and incubator settings were run at 37°C in 5% (v/v) CO_2_ in air.

**Figure 1:**
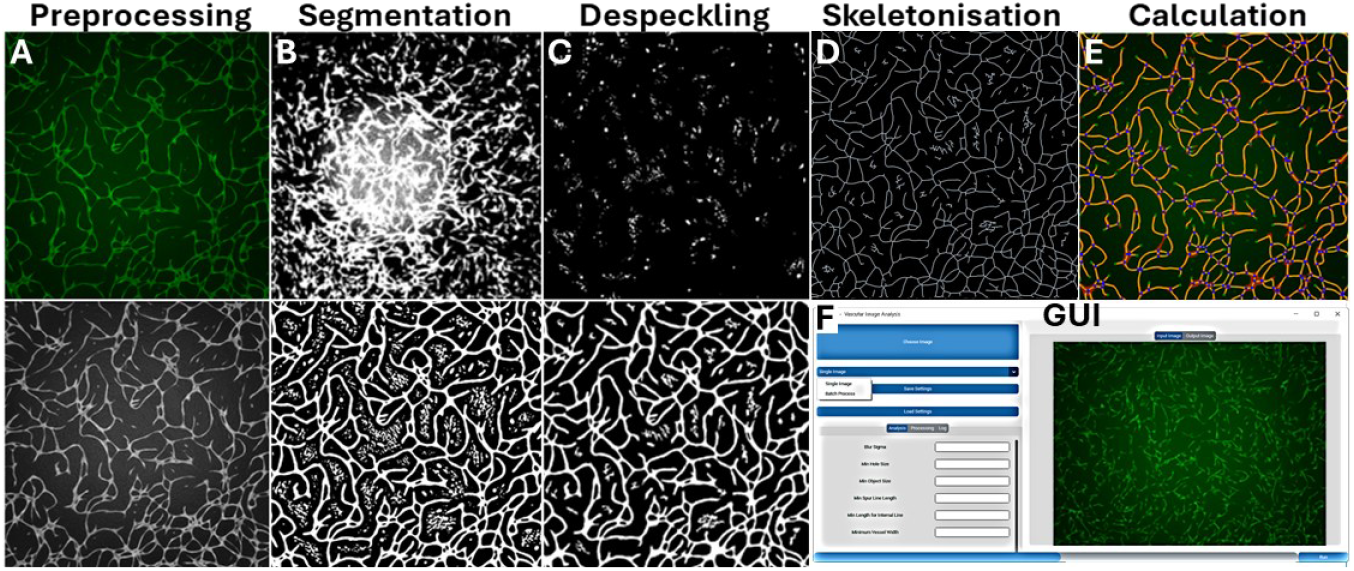
Illustration of BRAT process steps and improvements. (A) Vessel network fluorescence images were imported and relevant channels rendered in greyscale. (B) Segmentation was attempted by a global Otsu threshold value (top), then improved by individually thresholding local regions of the image to account for an uneven microscope illumination (bottom). (C) Segmented images were cleaned by removing small objects (speckles; top) resulting in a cleaned segmentation (bottom). (D) Skeletonization of the segmented mask was then performed requiring pruning of small skeleton branches or erroneously clustered branch node points, through a graph theory approach. (E) Resultant images of segmented microvessel shape, centreline, branches, and nodes were output, as well as metrics including total vessel area and length, average vessel diameter, and vessel branch point numbers. (F) Screen shot of BRAT’s executable software (.exe) graphic user interface (GUI). Further details of these steps can be found in supplemental figures S1-S5.

In **Fig. 1-3**, 2D vasculogenesis cultures were conducted following the optimised protocol of Murphy et al [29]. Briefly, green fluorescent protein-expressing human aortic endothelial cells (GFP-hAECs; TeloHAEC-GFP, ATCC) were routinely maintained in Vascular Cell Basal Media with Endothelial Cell Growth Kit-VEGF (#100030, 100041; ‘complete vascular media’) and human adipose-derived mesenchymal stem cells (hAD-MSCs; ASC52, American Type Culture Collection ATCC) in MSC Basal Media with Growth Kit Low Serum (#500030, 500040; ‘complete stromal media’). GFP-hAECs and hAD-MSCs were harvested from routine maintenance culture, counted via trypan blue exclusion method and mixed at a ratio of 1:5 cells (GFP-hAEC: hAD-MSC). This cell suspension was then centrifuged at 300g for 5 minutes, media aspirated, and cell pellet resuspended in complete vascular media. Cells were seeded at a density of 187,500 total cells/cm^2^ on clear flat-bottom polystyrene tissue culture-treated microplates and cultured for 7-days with complete media renewal on days 3 and 6. Culture plates were placed on the motorised stage of an in-incubator microscope (Etaluma LS720) and imaged by brightfield and GFP-hAEC fluorescence (473-491 nm excitation LED source, 502-561 nm detector) every 10 minutes from the 18^th^ to 165^th^ hour of culture (approximately 7 days; 886 timepoints). On days 3 and 6 of culture, cell culture media was exchanged as annotated by vertical lines in **Fig. 3**.

In **Fig. 5A**, 3D vasculogenesis cultures and microfluidic device preparation were conducted similar as the optimised protocol of Murphy et al [29]. Briefly, GFP-hAECs and hAD-MSCs were suspended in a 10 mg/mL fibrinogen solution in DPBS at 13×10^6^ cells/mL and 13×10^6^ cells/mL, respectively (26×10^6^ cells/mL total). This suspension was then mixed 1:1 with a solution of 4 U/mL thrombin in complete vascular media. Then 10 µL of this mixture was directly pipetted into custom fabricated microfluidic devices, previously described [29], at final concentrations: 13×10^6^ total cells/mL, 5 mg/mL fibrinogen, 2 U/mL thrombin. The cell-seeded device was then placed in a humidified incubator for 15 minutes to facilitate fibrin hydrogel formation. Media channels were loaded with complete vascular media and reservoirs filled with approximately 75 µL complete vascular cell medium each, and then cultured with complete media exchanges every 2–3 days for 7 days. GFP-hAECs in these 3D microchip co-cultures were live-imaged using the same Etaluma LS720 microscope setting and methods outlined above.

For **Fig. 5B**, Patient-derived cord blood outgrowth endothelial cells (BOECs, 6.5×10^5^ cells/mL) and hAD-MSCs (6.5×10^5^ cells/mL) were mixed into a solution of final concentrations 3 mg/mL fibrinogen, 1 U/mL thrombin, and 0.15 U/mL aprotinin, similar as above. A total of 30 µL of the mixture was placed in each well of 96-well plates and cultured in complete vascular media (200 µL) for 7 days with complete media exchanges every 2 days for 7 days. These BOEC and hAD-MSC 3D co-cultures were washed with phosphate buffered saline (PBS; 10010023, ThermoFisher) and fixed with 4% paraformaldehyde (C006, ProSciTech) in PBS. After three washes with PBS, co-cultures were permeabilised with 0.1% Triton X-100 (93443, Sigma-Aldrich) in PBS for 10 minutes and washed thrice with PBS again. Then, co-cultures were blocked with 10% goat serum (16210064, ThermoFisher) in PBS for 1 hour, stained with 2 µg/mL Mouse IgG2a anti-CD31 (MA3100, ThermoFisher) in 10% goat serum overnight at 4 ºC, washed thrice with PBS, stained by 4 µg/mL Alexa Fluor 555 Goat anti-Mouse IgG2a (A-21137, ThermoFisher) in 10% goat serum for 1h, and washed thrice with PBS. Finally, co-cultures were counterstained with DAPI (4′,6-diamidino-2-phenylindole, dihydrochloride, A-21137, ThermoFisher) for 10 minutes, washed thrice with PBS, and then imaged for CD31-positive microvascular networks on a Nikon ECLIPSE Ti2 Inverted Microscope (using a 555 nm excitation laser, 590-650 nm detector).

### 2.2. BRAT Image Processing

#### 2.2.1. Preprocessing

##### Import

The BRAT algorithm requires a folder location input, within which all compatible images will be processed. The Etaluma LS720 and REAVER dataset images used for this paper are TIFF file format. BRAT can also analyse JPG, PNG, and BMP formats. BRAT can directly import single-channel monochromatic images of vascular networks or can convert multi-channel RGB or immunofluorescent images to monochromatic images by taking an ‘individual channel’ selection, an ‘average of image levels’, or a ‘human eye level weighted average’ as described in its processing menu (**Fig. 1A**).

##### Intensity Rescaling

To normalize image brightness intensity and pattern across a batch of images, BRAT first rescales the brightest pixel in the image to full brightness and the darkest pixel to zero brightness. Then, BRAT applies a contrast-limited adaptive histogram equalisation to further increase the contrast of the image, allowing for small-local details to be visible even in regions that are darker or lighter than the rest of the image. This process is especially important given the large local differences in the lighting between different points in some images in fluorescence microscopy.

##### Blur Application

BRAT then applies a Gaussian Blur by convoluting a small Gaussian Kernel with the image to decrease small artefacts which may be present on the boundary of the future segmentation, as illustrated by **Fig. S1**. The size of this kernel is determined by the user; however, the performance addition of the blur is limited to a small kernel size. Larger sizes may cause an increased loss of contrast and with that the potential quality of the segmentation step.

#### 2.2.2. Segmentation

BRAT differentiates itself from other vascular network analysis tools by offering Local Otsu Adaptive segmentation, more robustly detecting vascular networks in images with variable background noise (**Fig. 1B, S2, S8**). Consistent with prior vascular network analysis tools, BRAT also offers traditional ‘Global Threshold (User Specified)’ and ‘Global Otsu’ segmentation options in its processing menu.

##### Traditional Segmentation Methods

During the development of BRAT, *global segmentation methods* were compared such as *Otsu-generated thresholds*.[30] This method finds a threshold value on which there is maximum inter-class variance and minimum intra-class variance. Each pixel is then compared against one global threshold value to find the segmentation, as implemented through Python’s SciKit-image morphology library.[31] Otsu segmentation is used by REAVER and many MVN tools, however, Otsu segmentation performed poorly in analysing the live-imaged vasculogenic assay, due to an uneven signal, perhaps from the laser or tissue culture plastic. Due to the uneven signal, the background in the centre of the image can be as bright as the foreground on the edges of the image (**Fig. 1B, S2, S8**).

##### Local Otsu Adaptive Segmentation

BRAT applies an averaging filter throughout the entire image, making a threshold image that has different threshold values throughout the image. The image is then compared to the generated threshold image to obtain the segmentation mask. BRAT’s averaging functions can be based upon the mean, median, and Gaussian weights, where Gaussian weights were chosen as the averaging function for this paper’s analysis, with the result shown in **Fig. 1B**.

This *region property-based local segmentation* using pixel brightness was chosen over *edge-based segmentation* and *active-contour methods*. Edge-based methods were rejected due to their unreliability in initial testing, as they created large numbers of false-region identification from the background.[32] In cases where an edge-detector does find a boundary of an image, the image contour must be closed for the region to be identified and added to the segmentation. Any holes in the identified boundary contour caused this issue to occur, further causing failures within an edge-based segmentation method.

*Active-contour-based segmentation methods* were also considered for the segmentation,[33] but due to the high number of potential individual regions to be segmented, the number of iterations required to get a segmentation was considered unacceptable. This problem was most present during the beginning and middle of the vasculogenesis culture, where there were large numbers of individual regions to be segmented. At the end of these cultures, the cells within these images had formed MVNs but presented issues in relation to the hollow shapes of the MVN, meaning that inside contours also had to be generated for accurate segmentation, further increasing the number of iterations required. Finally, other methods were attempted, including a method based on *k-means segmentation*, but were not useful here.

##### Segmentation Cleaning

Post-segmentation, vessel areas may include holes within the vessels as well as small groups of cells. These may be considered undesirable to researchers and as such should be removed. Within the config file of the system, a researcher can set minimum hole and object feature size that can be removed automatically by the program from the segmentation (similar to AngioTool). These are not included in any of the metrics generated from the segmentation or its children. This process is illustrated in **Fig. 1B** and **Fig. 1C**, with original MVN segmentation, hole filling at 300 pixels, and minimum size threshold at 400 pixels.

#### 2.2.3. Skeletonisation

Skeletonisation was performed by successive erosion of the segmented MVN through a medial-axis transform using the SciKit image morphology library (**Fig. 1D**).[31] However, this transform produces spurs in the MVN skeleton that reflect the positions where the vessel may not be strictly symmetric or otherwise has some level of outcropping from the parent vessel.

##### Vessel Width Pruning

An option for pruning the skeleton is to consider the width of the vessel. This value can be set in the configuration file by the user to control the minimum width in terms its width. This is implemented by calculating the euclidean-distance transform (EDT) of the segmentation and applying the original skeletonisation as a mask upon it. This leaves a skeletonisation which instead of being a binary image has its EDT encoded within itself. This skeletonisation can then have a threshold applied to find positions that have the minimum line width, as at the bottom of **Fig. 1D**.

#### 2.2.4. Graph Processing

Graph-based processing is a novel MVN skeletonization approach, producing a mathematical structure consisting of individual nodes connected to each other with edges. These edges can have weights or directionality associated with them and in these cases, they are termed weighted and directed graphs respectively. An MVN’s skeleton image can be represented as a weighted, undirected graph. A node on the graph represents the junction and end points of the skeleton while the edges represent the connections between them. The weights represent the distance to traverse along these edges. These were implemented using the Python library, *NetworkX*.[34] It can be noted that a list of pixel positions can be stored in each edge, meaning that from the graph data structure, the entire skeleton can be reconstructed. The graph representation of the skeleton has a high initial computational cost due to the slow, recursive nature of the initial graph construction. Graph representations are advantageous in terms of the ease of processing MVN skeletons for measurement, and of pruning or similar processes, in comparison to 2D image topology with larger matrices and computational costs. This is an innovation for MVN analysis tools, allowing for more efficient and new computational processes to analyse MVN skeletons.

##### Graph-Based Length Pruning

Length based pruning is the process of removing spurs on the skeletonisation based upon their length. If the length of the vessel is found to be less than that of a threshold set by the user in the user configuration file, then the vessel will be removed. The method of operation for this process is based upon the process of determining the length of each spur point. If a skeleton splits into multiple points, the longest of the spurs is considered as the continuation of the original skeleton portion while other lengths are considered as spurs. An algorithm process diagram and illustrations for this pruning vessel edges can be found in **Fig. S3, S4**.

##### Node Consolidation

Node consolidation is a process by which nodes that are deemed sufficiently close together can be consolidated into a single junction point. This process works on internal nodes that are connected and if they are below an independent user-set threshold, the two separate nodes are consolidated to a single node with the internal edge between them being split in half and its weight and points added to the edges that connected to the original nodes. An algorithm process diagram and illustrations for this technique can be found in **Fig. S3, S5**.

##### Skeleton Reconstruction

Following the aforementioned graph-based modifications, the skeleton must be reconstructed. It can be noted that as part of the original conversion that each edge stored a list of all points within the image that corresponded with that edge. With this inclusion to the data structure, it is trivial to reconstruct the skeleton by simply looping over each edge within the graph and setting all points stored by the skeleton edge to ‘True’ in the binary image. An example skeleton overlaid on a segmented vessel can be found in **Fig. 1E**.

### 2.3. Implementation and Measurement

#### Implementation

BRAT’s executable software interface allows users to input process parameters such as: blur sigma, minimum hole size, minimum object size, minimum spur line length, minimum length for internal lines, minimum vessel width, and an image directory path for single-image or entire-folder processing modes (**Fig. 1F**). BRAT creates a list of all images within the path’s list which can then be imported to the system. The algorithm processes are applied to each image in order of its position on the list, whose results are stored as a Python Pandas DataFrame and exported to a .csv file to a user-specified location. Implementation speed was tested on a typical laptop (CPU: Intel Core i7-1065G7, RAM: 16 GB, GPU: Intel Iris Graphics) and a high-performance computer (HPC; CPU: AMD Ryzen Threadripper PRO W955WX, RAM: 128 GB, GPU: Nvidia RTX A5000)

#### CPU Multi-Threading

BRAT’s batch image processing takes advantage of computer processor multi-threading, allowing it to analyse several images in parallel, requiring a fraction of time as other existing image analysis algorithms. Multi-threading is implemented using the Python MultiProcessing library.

#### Total Segmented Area

The segmented area is calculated using only the segmentation mask image. It simply consists of counting all non-zero pixels within the segmentation mask image, as in **Fig. 1E**.

#### Total Vessel Length

The total vessel length is calculated similarly to that of the total segmented area. The number of non-zero pixels are counted within the skeleton image.

#### Number of Vessels and Average Vessel Length

The number of vessels is determined by counting the number of edges within the Graph MVN representation. Each of the edges’ weights are removed from the list of graph edges and the mean is taken.

#### End Points and Junctions

The number of End Points and Junctions are determined by counting the number of nodes with only one connected neighbour as well as the number of nodes with greater than one connected neighbour within the Graph MVN representation.

#### Average Vessel Width

The Euclidean distance transform (EDT) was computed to allow vessel width tuning to take place. The EDT was masked by the skeleton image to obtain an image/matrix that described the half-width of the vessels at every point on the skeleton. To calculate average vessel width, the mean of all non-zero values within this image matrix.

## 3. Results and Discussion

### 3.1. BRAT is the fastest batch analysis tool of high-content microvascular network datasets

Vasculogenesis culture assays were performed by co-culturing GFP-hAECs with hAD-MSCs in complete vascular media, as in Section 2.1. Briefly, culture images were captured every 10 minutes from hour 18 to hour 165 of culture (7 days, 886 timepoints) by an in-incubator microscope at 4x magnification. The green fluorescence images of GFP-hAEC microvascular networks were batch analysed under identical parameters by BRAT. Every tenth image (89 images captured every 1.6 hours) was analysed using AngioTool, where an operator spent additional time to manually select optimal parameters for each image, shown in **Fig. 1, 2, 3, S7, S8**.

**Figure 2:**
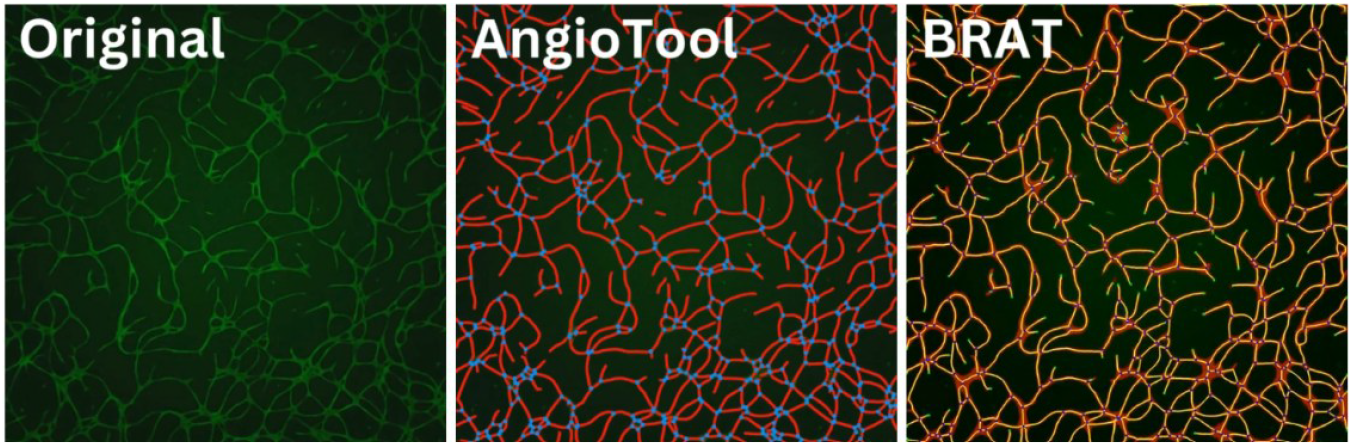
Real-Time video microscopy of an in vitro vasculogenesis assay, with BRAT and AngioTool outputs. Co-cultured GFP-hAECs and hAD-MSCs form microvascular networks over 7 days, as imaged every 10 minutes. BRAT analysed all 886 timesteps, while AngioTool only analysed 89 timesteps. In the bioRxiv version, the video is a download in the ‘Supplemental Material’ tab on the right of the abstract.

**Figure 3:**
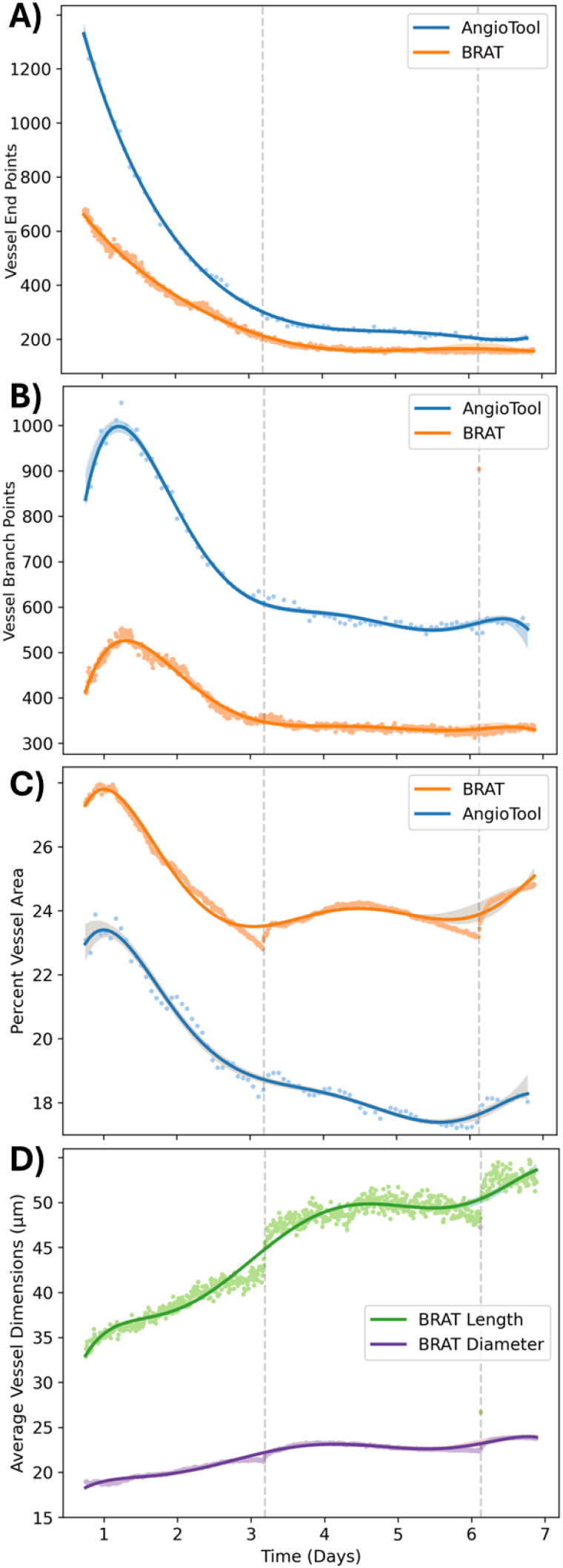
Batch-processing of a live-imaged vasculogenesis assay. GFP-hAECs co-cultured with hAD-MSCs for 7 days to form microvascular networks, with images captured every 10 minutes, as Fig 2. BRAT and AngioTool were applied to measure microvascular network (A) end points, (B) branch points, and (C) vessel area as a percentage of the field of view area. (D) BRAT can also measure average or total vessel diameter and length (distances between end or branch points). Hashed vertical lines represent media change times. Individual timepoint analysis (dots) are overlaid with a regression and 95% confidence intervals. These results are compared with REAVER in **Fig. S8.**

The live-imaged vasculogenic assay starts by imaging single GFP-hAECs; analysed by BRAT and AngioTool as a microvascular network (MVN) with many end points (**Fig. 2A**) and few branch points (**Fig. 2B**) of moderate MVN area (**Fig. 2C**) yet very small vessel segment length and diameter (**Fig. 2D**). Consistent with our previously reported observations [29], the GFP-hAECs proliferate and form small discrete vessels from days 0 to 1, which decreases end point and increases branch point numbers, and comprise larger MVN lengths, diameters, and total area. From day 1, the MVN condenses in end points, branch points, and area but with increasing vessel length and diameter, and this MVN condensation seems to stabilize by day 4.

**Fig. 2, 3, S7**, and **S8** compare BRAT and AngioTool’s ability to analyse this live-imaged MVN assay. AngioTool seems more sensitive than BRAT in identifying isolated cells (end points) and finely segmenting microvessel network branch points to correctly identify small holes, while BRAT’s segments a consistently larger MVN area. AngioTool’s sensitive segmentation sometimes leads to false-positive end and branch points (**Fig. S7**). It can be noted that there is more variation in the end point count made by BRAT, potentially due to pixel speckling, which could be mitigated by increasing the minimum vessel area. In BRAT and AngioTool, increased MVN area, diameter, and length occurred directly after media feeding on days 3 and 6. This could be a result of increased cell growth or simply altered condensation on the tissue culture plastic surface after feeding.

AngioTool requires 70-fold more processing time than BRAT on an HPC, mostly direct operator input. When processing 89 images of our 7-day vasculogenesis assay, AngioTool requires the user’s attention for 16 seconds per image to manually adjust software tools. In contrast, BRAT requires the user to identify process parameters once, and then all 886 images can be processed at 0.17 seconds/image on a HPC or 2.31 seconds/image on a typical laptop. Multi-threading allowed images to be processed 8 times faster on the HPC and 4 times faster on the laptop. The most computationally intense functions were shown to be the draw and save algorithm (3.9 seconds) and the skeletonisation algorithm (3.2 seconds), accounting for 85% of the total run time (**Fig. S6**). AngioTool is considered a leading MVN analysis algorithm, in the next section our benchmark comparisons using REAVER’s broad MVN image dataset indicate that BRAT provides a higher quality MVN segmentation and analysis than AngioTool.

REAVER is regarded as, to date, the most accurate open-source MVN analysis for immunofluorescent histology.[17] In **Fig. S7, S8**, we assess REAVER’s ability to semi-automatically segment the live-imaged vasculogenesis assay. REAVER required an average of 13.7 seconds to analyse one image on a HPC, or 20:32 to analyse only 89 images (**Fig. S7**). Despite its rapid skeletonization (3.2s/image) and MVN metrics (2.8s/image), 12.6s was required by the REAVER software for a user to perform a manual inspection of each image to tick an ‘output’ checkbox; often delayed by GUI errors for our large dataset. REAVER’s segmentation of the live-imaged dataset appear poorer than both BRAT and AngioTool (**Fig. 2, S7, S8**), unable to detect 30% of MVN segments. This appeared to result from REAVER’s global thresholding *vs* BRAT’s local methods (**Fig. 1B**), which created background noise artefacts in REAVER. Therefore, REAVER’s segmentation improved when thresholds were manually adjusted for each time point (**Fig. S8**), at the consequence of losing automation and high-throughput screening time.

### 3.2. BRAT is the most sensitive microvascular network segmentation tool

The performance of BRAT was evaluated against four leading microvascular network (MVN) image analysis algorithms: AngioQuant, AngioTool, RAVE, and REAVER, using 36 images of microvascular networks of diverse sources and tissues previously provided as a quality control metric by REAVER.[17] Accuracy (*S*^*A*^), sensitivity (*S*^*N*^), and specificity (*S*^*C*^) can be calculated as the number of true positive (TP), true negative (TN), false positive (FP), and false negative (FN) detections of pixels or nodes as compared to manual scoring, by the following equations:

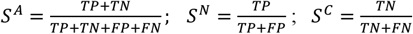

However, AngioTool does not return segmented images to measure *S*^*A*^, *S*^*N*^, and *S*^*C*^ at pixel resolution (**Fig. 4A**) or provide vessel diameter or length (**Fig. 4B**). REAVER measured AngioTool outputs using a screenshot of AngioTool’s GUI during segmentation, then processed by REAVER’s output metrics.[17]

**Figure 4:**
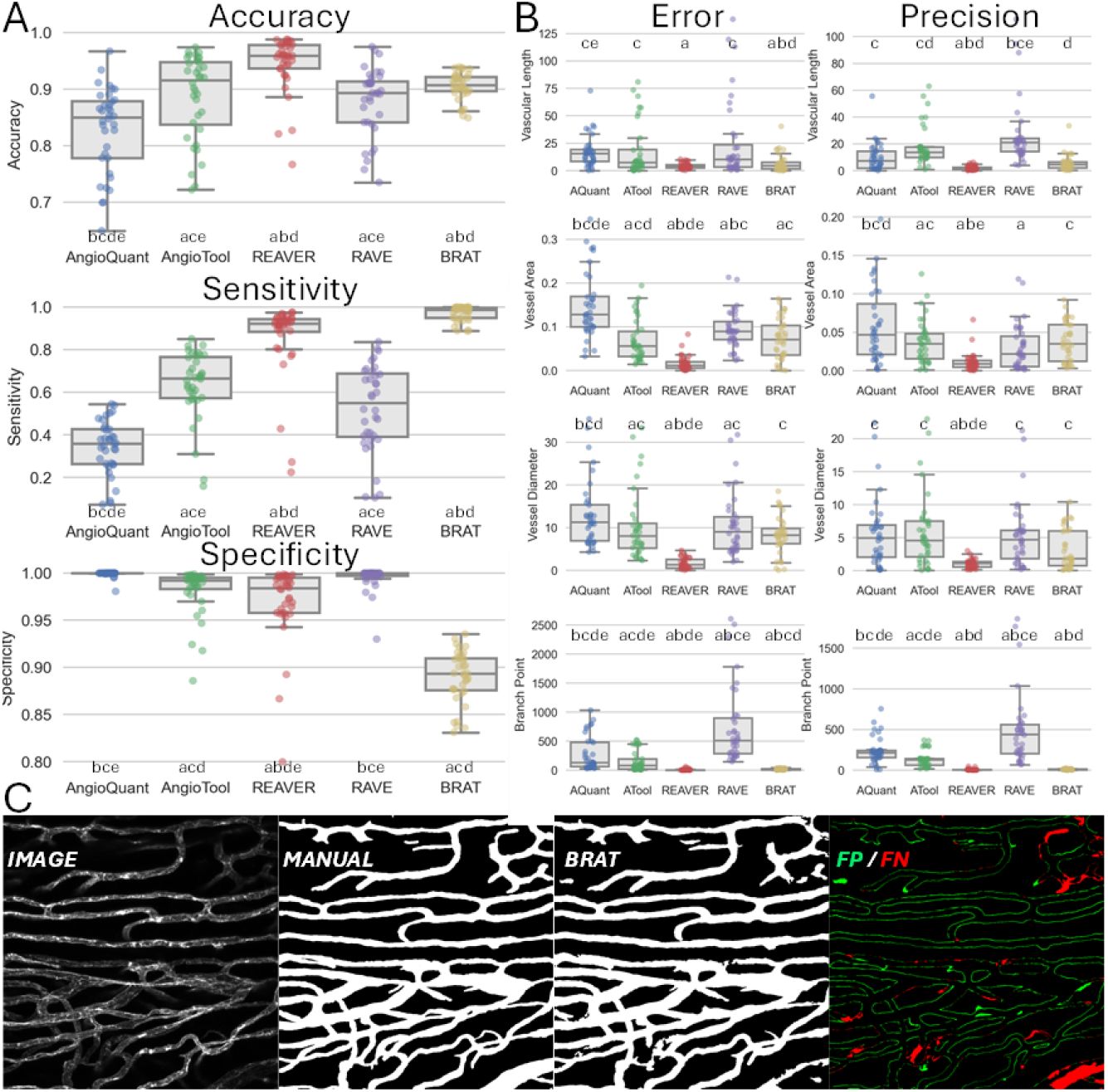
BRAT benchmarked against vessel analysis software using a microvascular image database. (A) A comparison of BRAT versus the leading four open-source software’s accuracy, sensitivity (true-positive), and specificity (true-negative) of microvascular network (MVN) pixel segmentation versus gold-standard manual segmentation. (B) Accuracy and precision across automatically detected vessel length, area, diameter, and branch points with reference to manual segmentations. (C) The most challenging MVN image to segment from the REAVER dataset; an image of CD31^+^ MVN in cardiac tissue (left), beside a manual annotation (ground truth), BRAT’s automated annotation, and BRAT’s false-positive (FP; green) and false-negative (FN; red) incorrectly-labelled MVN pixels. (A, B) Boxplots represent mean, quartile ranges, while whiskers extend to show the distribution, less outlier points. The annotations above or below each plot indicate significant pairwise comparisons between groups with Bonferroni adjusted p-values (letters).

In **Fig. 4A**, BRAT’s accuracy (93.3% ± 0.3%) is second only to REAVER (94.4% ± 0.5%), and BRAT was the most consistently accurate segmentation tool with the tightest range of accuracy measurements. BRAT’s sensitivity (94.5% ± 0.6%) performs better in direct comparison with any other tool (*p* < 0.013), implying BRAT is the best tool to segment regions known to be a part of MVN. However, BRAT is the worst tool for specificity (93.6% ± 0.2%) a result which could be biased by errors in under-segmented ground truth images (**Fig. 4C**). Additionally, tools with the highest specificity, namely AngioQuant and RAVE, also exhibit the lowest accuracy. Therefore, the relative performance in this metric may not be a result of the quality of the tool’s segmentation but more to do with a tool’s failure to segment MVNs.

Metrics relating to the tool measurement error (*E*_*tool*_) and precision (*P*_*tool*_) are shown in **Fig. 4B**. Error is defined as the absolute difference between the tool’s measurements (*Y*_*tool*_) and manual measurements (*Y*_*manual*_) for 36 images in the REAVER publication (immunofluorescence images of microvascular networks in diaphragm, heart, adipose tissue, liver, peritoneal cavity, retina) [17]. Precision is defined as the absolute difference in each image’s measurement error (*E*_*tool*_) to its average error (*E*_*tool*_).

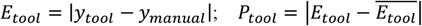

BRAT performed at a similar quality or better than that of AngioTool in measuring vessel length, vessel diameter, and branch point numbers (p < 10^−4^). In most metrics, the error and precision of BRAT exceed that of AngioTool, and all other tools excluding REAVER.

To determine the source of BRAT’s low specificity and errors in measuring vessel length, in **Fig. 4C** we consider the image that had the worst specificity in the REAVER Dataset, a 20x magnification of a capillary network from heart tissue. RAVE’s analysis of this image measured microvascular networks with 94% specificity as compared to manual segmentation, its worst measurement. Its segmentation is difficult due to the high variation in pixel brightness between different, closely placed vessels. To determine the failure within the segmentation, **Fig. 4C** shows an operator’s hand-selected MVN pixels considered as ‘ground truth’, in comparison to BRAT’s automatically segmented MVN pixels. The False Positive pixels produced by BRAT are also included. Comparing the locations of the error points to that of the original image in **Fig. 3C**, some of BRAT’s false positive pixels appear to be part of the MVN, while some manually segmented pixels do not. Reviewing similar images from the REAVER dataset, we do not believe BRAT’s lower specificity on REAVER’s dataset images to be a critical limitation.

### 3.3. BRAT can analyse diverse 2D and 3D Microvascular Network Datasets

To further demonstrate the broad applicability of BRAT, we applied it to analyse other microvascular culture images, more complex than 2D GFP-hAEC:hAD-MSC vasculogenic culture images (**Fig. 3**) or REAVER’s animal and human histology slides (**Fig. 4**). In **Fig. 5** we demonstrate BRAT’s ability to analyse more challenging 3D microvascular networks from 2D fluorescence images, including a GFP-hAEC:hAD-MSC co-culture and a human umbilical cord blood outgrowth endothelial cell (BOEC)-hAD-MSC co-culture embedded in a 3D fibrin hydrogel (**Fig. 5A, B**). The 3D BOEC:hAD-MSC co-cultures were analysed to have many vessels of significantly smaller length than either 2D or 3D GFP-hAEC:hAD-MSC co-cultures (**Fig. 5C**). In contrast, 3D GFP-hAEC:hAD-MSC co-cultures had significantly thicker vascular network diameters and fewer branch points than other cultures. The 3D co-cultures exhibited higher vessel coverage area, a 2D image calculation biased due to the overlapping nature 3D microvascular networks in the 3D hydrogels. Additional BRAT metrics are shown in **Fig. S9**.

**Figure 5:**
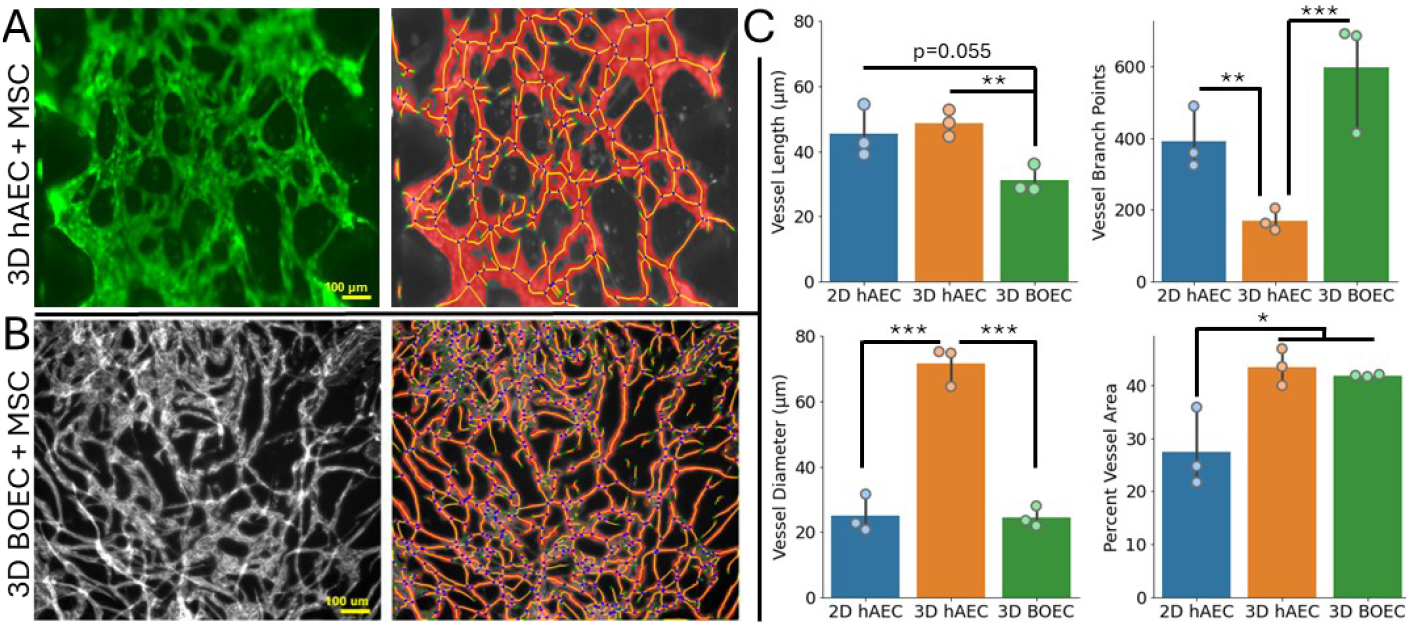
BRAT characterises diverse microvascular network images. (A) 3D microvascular networks formed by GFP-hAECs and hAD-MSCs co-cultured in a hydrogel microchip. (B) A 3D microvascular network of CD31-stained umbilical cord blood-outgrowth ECs (BOECs) and hAD-MSCs co-cultured in a fibrin hydrogel. (C) BRAT shows hAECs form thicker, longer, but fewer microvessels in 3D hAD-MSC hydrogel co-culture as opposed to BOECs. (A, B) 100 µm scale bars. (C)’s dimensions are reported in µm using 0.81 µm/pixel (Etaluma 720S) or 0.73 µm/pixel (Nikon Ti2) conversion factors. (C) datapoints represent independent cultures. Barplots represent mean ± standard deviation. *p<0.05, **p<0.01, ***p<0.001. Additional BRAT-generated metrics can be found in Fig. S9.

## 4. Conclusion

In this paper, we create and introduce the Batch-Resourcing Angiogenesis Tool (BRAT) as a state-of-the-art computer algorithm and software for the high-content screening of imaged microvascular networks (MVNs). BRAT follows a traditional image analysis process of preprocessing, segmentation, skeletonization, and characterisation. BRAT uses local signal normalisation to enhance segmentation quality and consistency and graph network representation to improve skeleton processing and analytical speed (**Fig. 1, S2**) an advantage over existing global segmentation tools (Section 2.2.2, **Fig. S7, S8)**.

Compared to all existing vascular network image analysis tools, BRAT exhibits the leading sensitivity in network segmentation as well as leading accuracy across a variety of metrics, including vessel area, diameter, and branch point number (**Fig. 4**). Compared to the REAVER image dataset benchmark [17], BRAT exhibits slightly lower accuracy than the REAVER tool due to a lower specificity (false positive), leading to erroneously high vessel length, area, and diameter (**Fig. 4B**). BRAT outperforms the REAVER tool on this paper’s live-imaged vasculogenesis assay (**Fig. 2, 3, S7, S8**) and in reviewing BRAT’s performance on REAVER’s manually-scored dataset (**Fig. 4C**), we posit low BRAT specificity is due to an overly-conservative manual segmentation.

BRAT is the only fully automated open-source MVN segmentation tool, since REAVER and other tools require time-significant manual interventions to analyse each image. BRAT’s capability to rapidly batch process thousands of MVN images (**Fig. 2, 3**) at 10-to-100-fold faster computational times (sub-seconds per-image) and without user input makes BRAT a viable tool for high content screening of angiogenic networks in live cell culture imaging and high-throughput drug screening. BRAT’s ability to analyse diverse types of MVN cell cultures includes 3D hydrogel co-cultures of hAD-MSCs with human aortic endothelial cells (hAECs) or umbilical cord blood outgrowth endothelial cells (BOECs) (**Fig. 5**). BRAT precisely detects statistical differences in these MVN geometries from only 3 culture replicates.

Microvascular network analysis is of increasing importance in diagnostic medicine, preclinical (animal) trials for new drugs, and for in vitro (cell culture) screening of thousands of drug candidates at different concentrations. Clinical indications with diagnostic analysis of microvascular networks include cardiac or cerebral ischemia [35], diabetic revascularisation [36], tumour vascularisation [13,14], and diverse microangiopathy in response to disease, such as COVID-19 [37]. The BRAT software presented in this paper not only achieves leading MVN segmentation and measurement sensitivity but importantly delivers a tool able to consistently analyse massive datasets of images with minimum operator input. BRAT’s image analysis time (sub-seconds per image, prior tools are tens of seconds per image), and BRAT’s standardised analysis will support the translation of high-content cardiovascular drug screening studies where microvascular network formation is a readout.

## Acknowledgements

This work was supported by Advance Queensland Industry Research Fellowships (AQIRF1312018), Australian Research Council Fellowships (DE220100757), and Ramaciotti Health Investment Grants (2022HIG08) to MCA. This work used the Queensland node of the NCRIS-enabled Australian National Fabrication Facility (ANFF) to fabricate silicone microchips for hydrogel cultures.

## Statements and Declarations

### Contributor Role Taxonomy (CRediT) Statement

**HK**. Harriet programmed all code and software, preformed timecourse and benchmark analyses, and helped write the first draft of the paper: conceptualization, formal analysis, investigation, methodology, software, validation, visualization, writing – original draft.

**ARM**. Ash co-supervised Harriet, preformed cell cultures, ran software analyses, and revised the paper: formal analysis, investigation, methodology, supervision, writing – review & editing.

**RM**. Ryan preformed cell culture, ran software analyses, and revised the paper: formal analysis, investigation, methodology, writing – review & editing.

**RAF**. Rose Ann preformed cell culture, ran software analyses, and revised the paper: formal analysis, investigation, methodology, writing – review & editing.

**MCA**. Mark was the main supervisor, ran software analyses, designed figures, helped write the first draft of the paper, and revised all drafts. Conceptualization, formal analysis, funding acquisition, project administration, supervision, visualization, writing – original draft, writing – review & editing.

## Supplementary Information

**Fig. S1:**
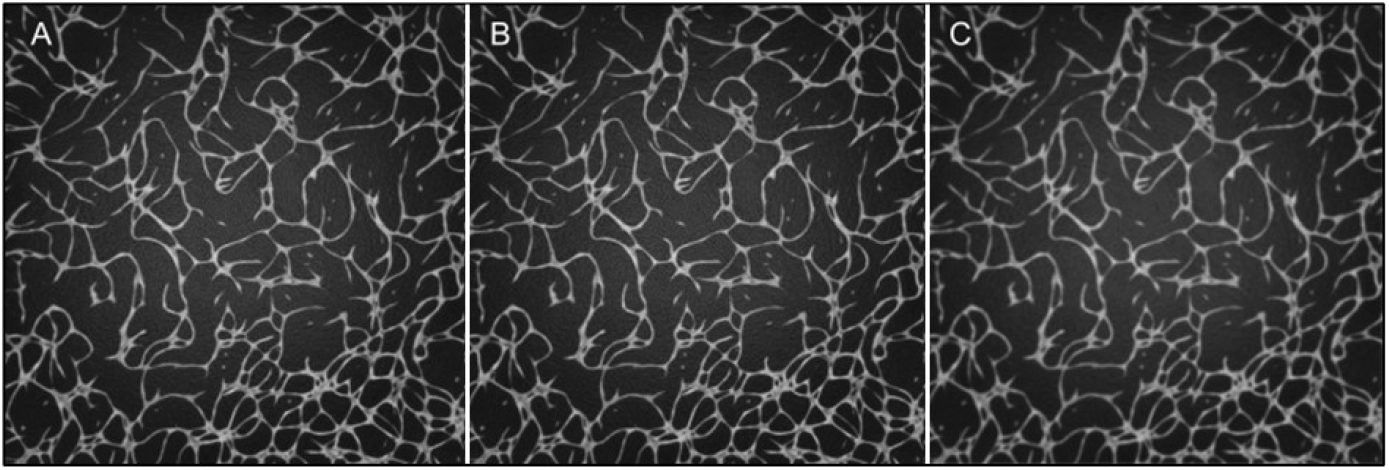
Application of Gaussian blur processes to a MVN image. (A) Reference image. (B) σ = 0.5 Gaussian blur. (C) σ = 3 Gaussian blur.

**Fig. S2:**
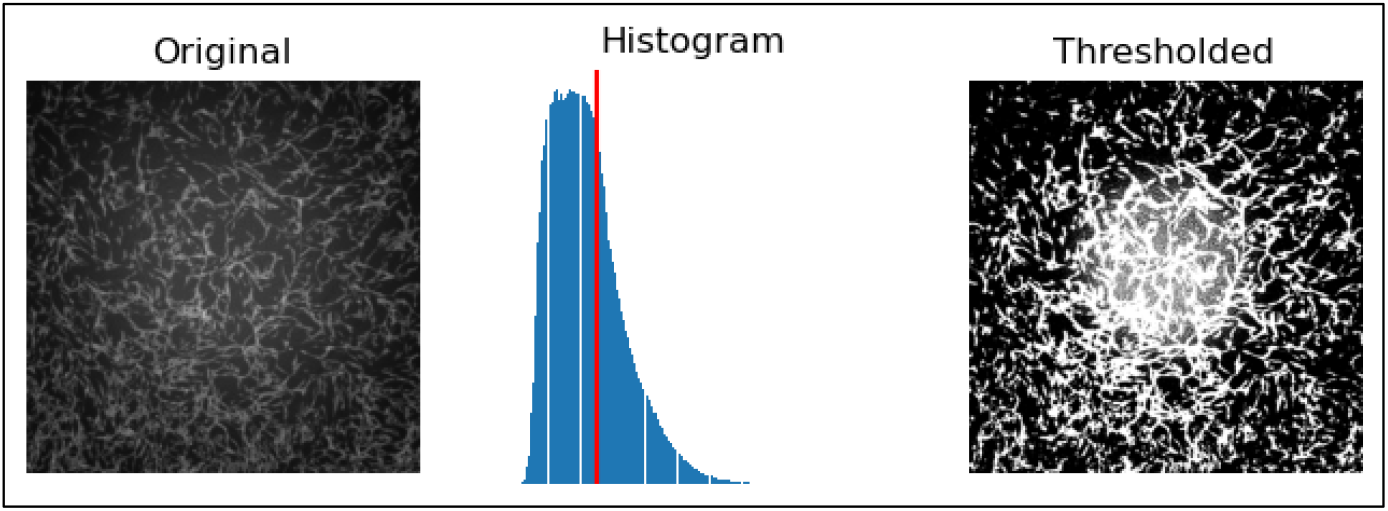
Illustration of poor performance of MVN segmentation using a global Otsu segmentation. Original GFP-hAEC fluorescence image on left, Otsu-calculated threshold in centre, and segmentation on right showcasing a central laser light artefact with increased brightness in the image centre.

**Fig. S3:**
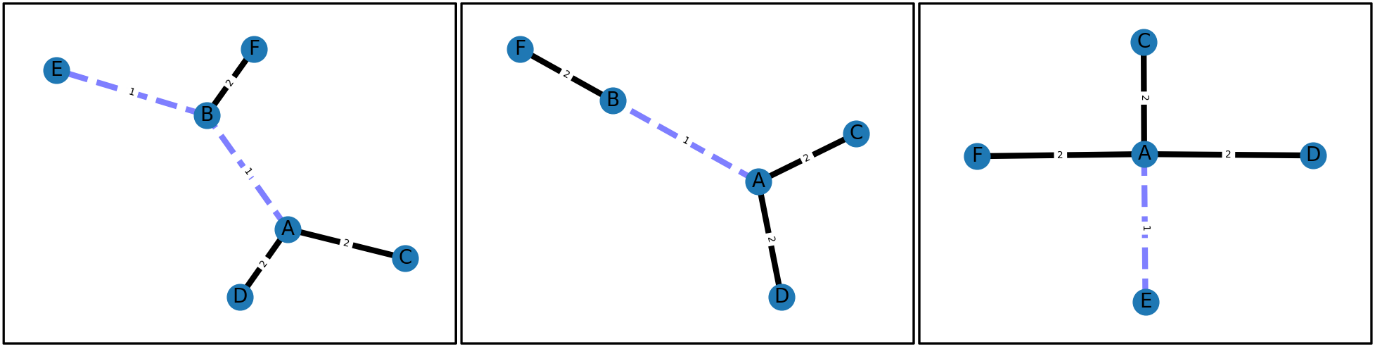
An illustration of Graph Network Representation of MVN (left), after removal of short spur edge E-B (centre), and after consolidation of small internal edge A-B (right).

**Fig. S4:**
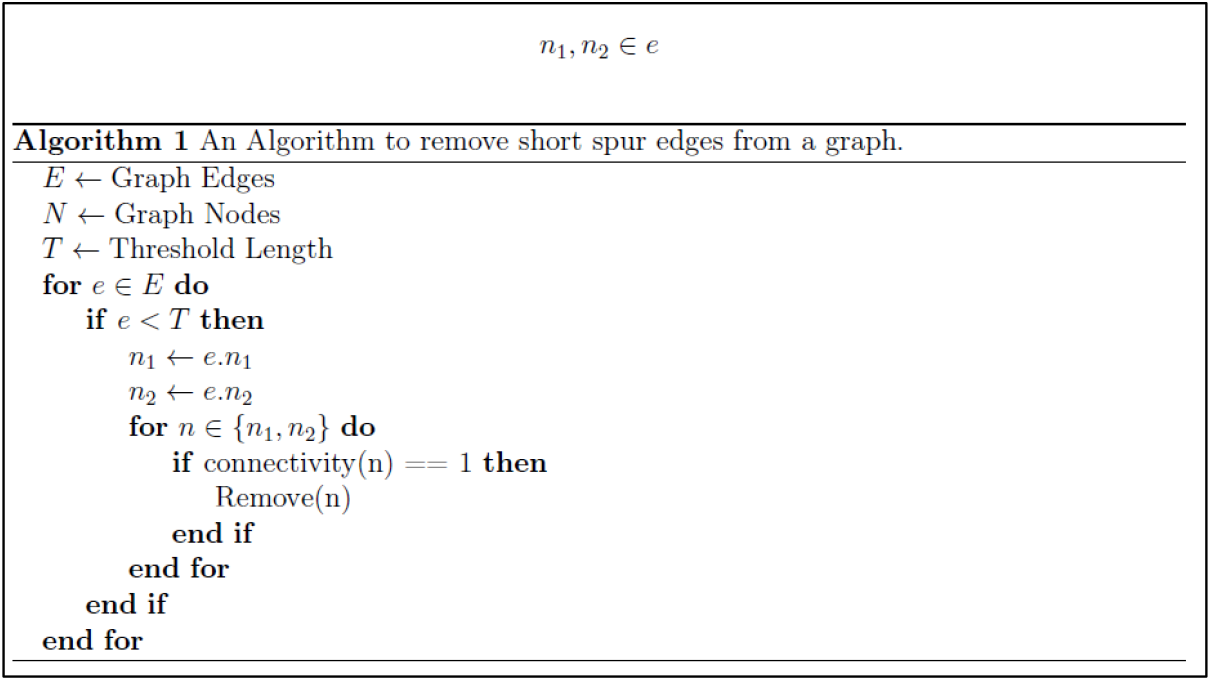
An algorithm to remove short spur edges from a MVN using Graph Network Representation.

**Fig. S5:**
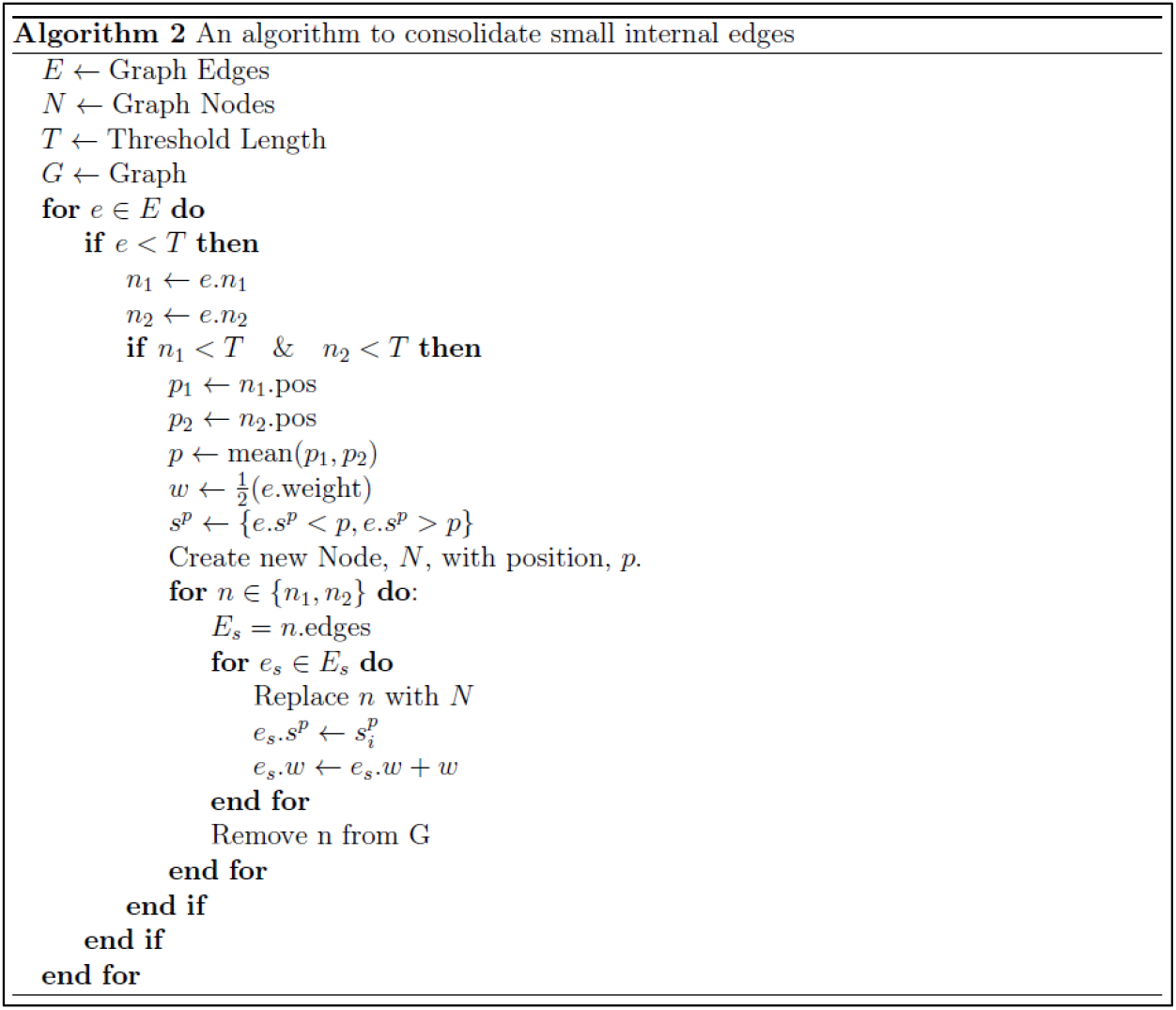
An algorithm to consolidate small internal edges from a MVN using Graph Network Representation.

**Fig. S6:**
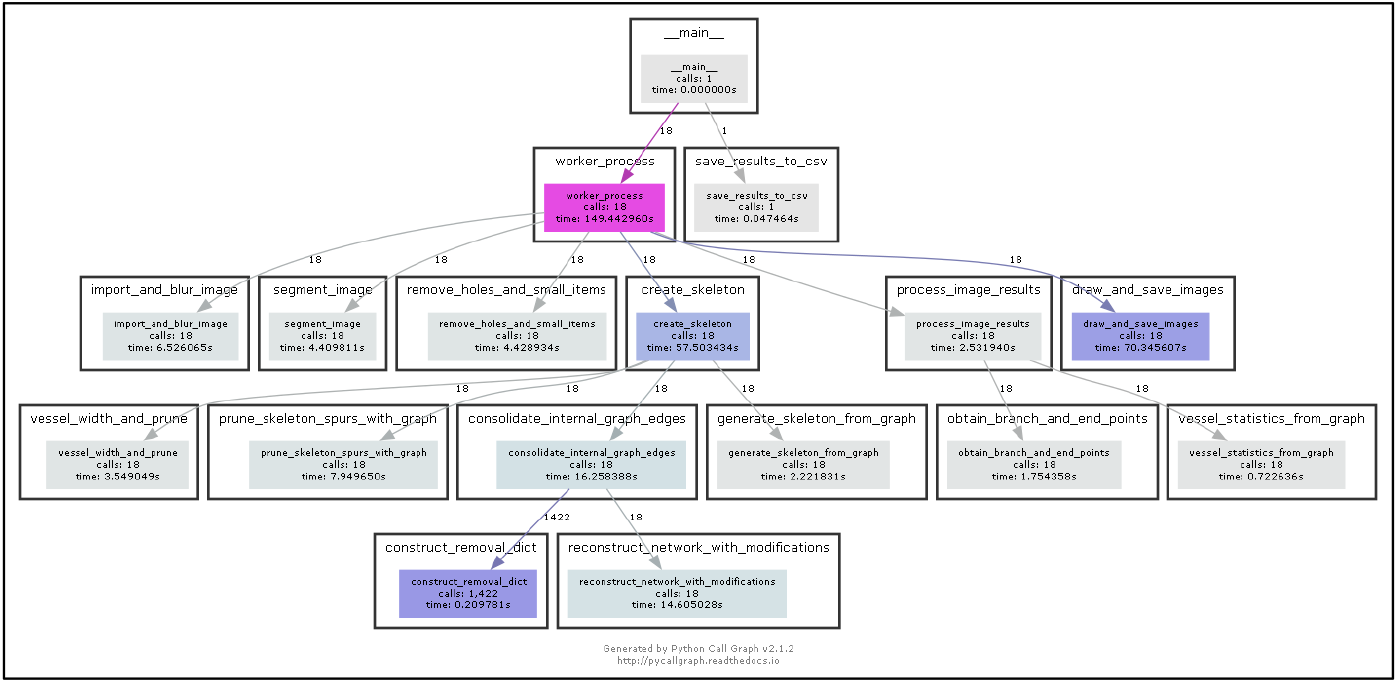
BRAT process call graph with process computational times for the HPC (N = 17 images).

**Fig. S7:**
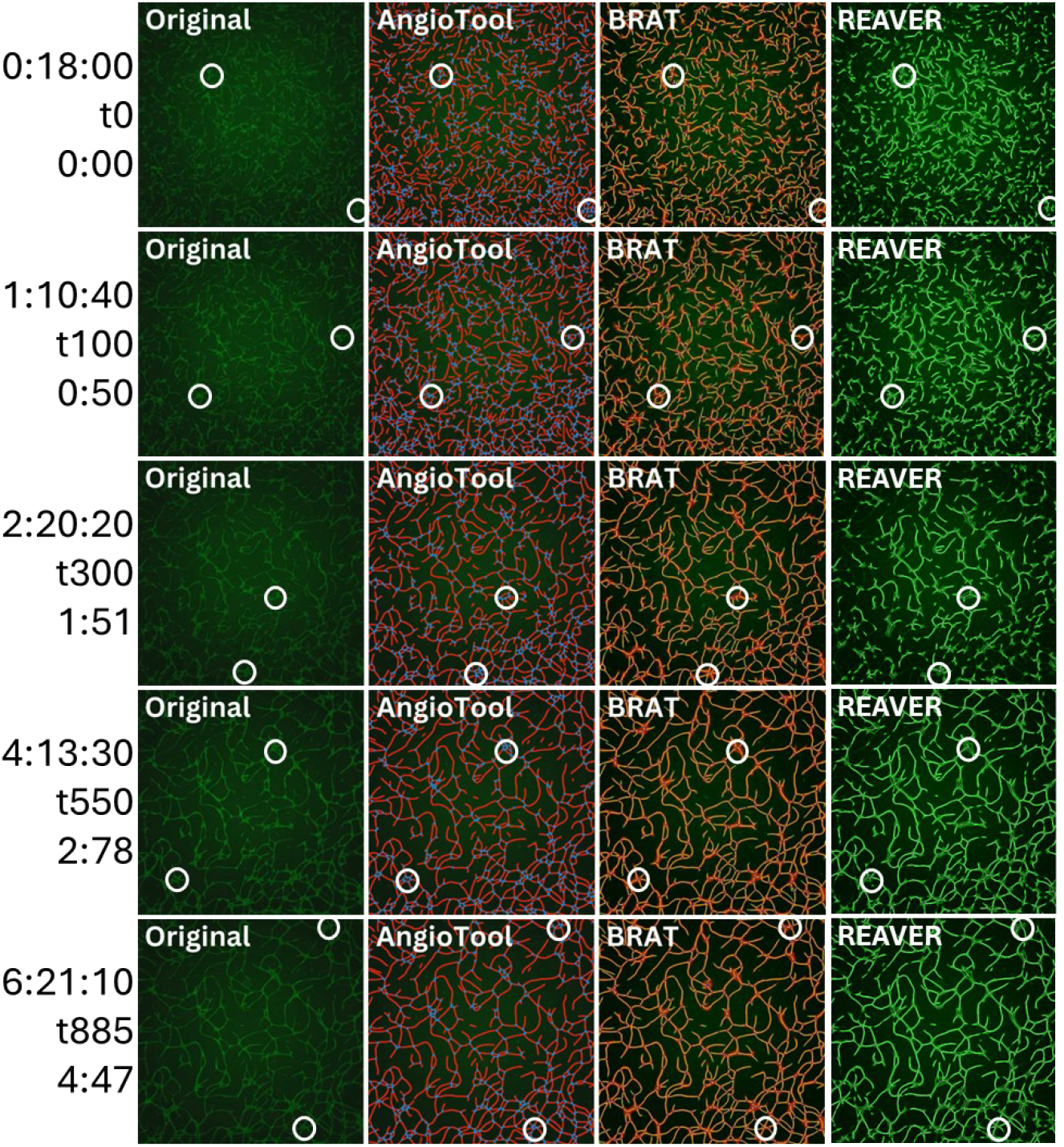
Visual comparison of vasculogenic images compared to AngioTool, BRAT, and REAVER segmentations. Three time annotations are on the left hand side: (top) d:hh:mm into cell culture similar as **Fig. 3**, (centre) timepoint number [0-885], (bottom) video timestep corresponding to **Fig. 1** (m:ss). Two subregions are circled in each image; one where AngioTool and one where BRAT exhibited errors. REAVER images are duplicate as in **Fig. S8A**, analysed with the threshold parameter set to 0.09.

**Fig. S8:**
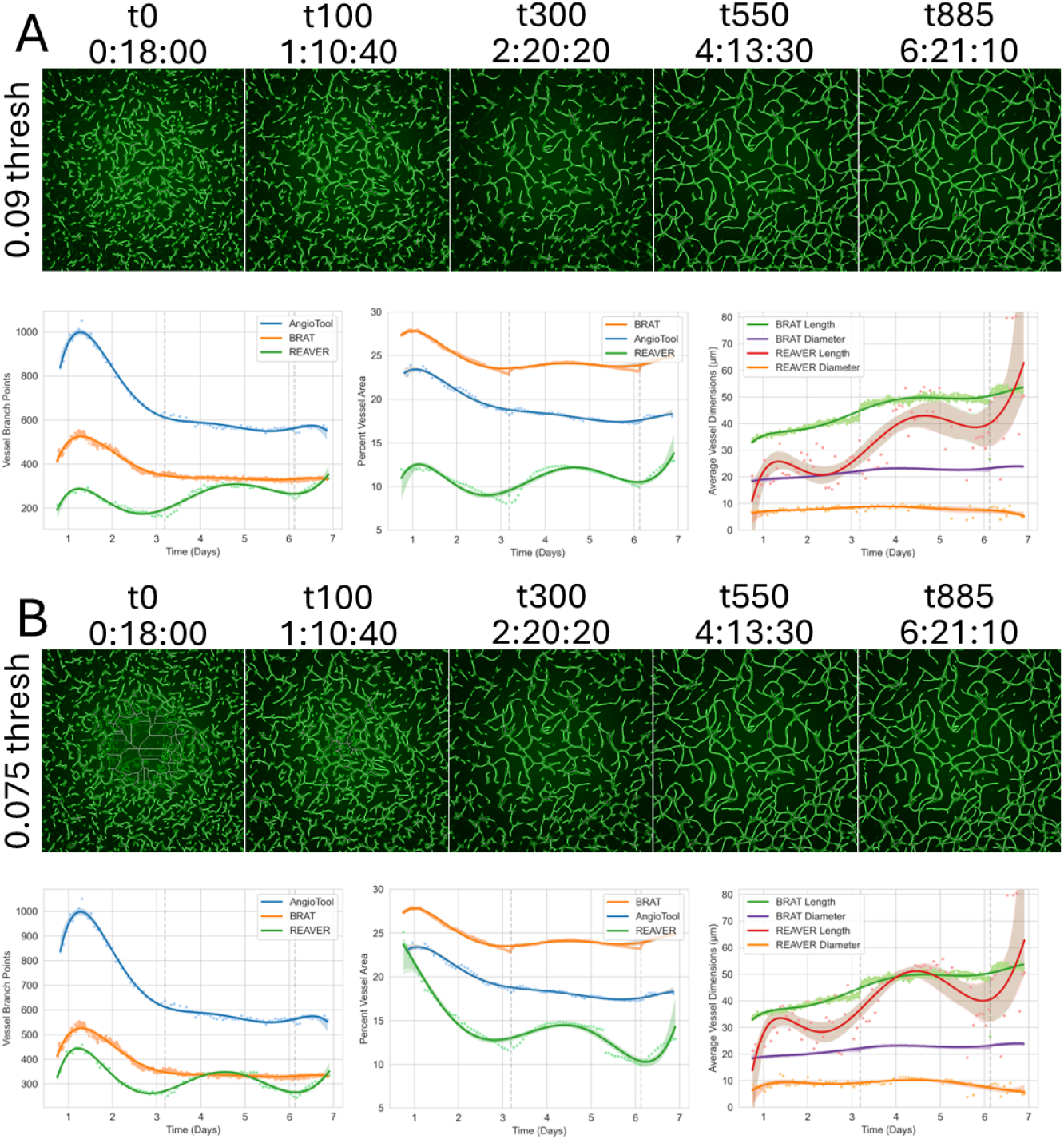
Comparing REAVER with AngioTool and BRAT for time-lapse imaging of microvascular network formation. Threshold at (A) 0.09 (t0-optimal) or (B) 0.075 (t300-optimal). REAVER Settings: Res. 0.8125, Avg Filt Size: 128, Min Conn-Comp A: 50, Wire Dil Thresh: 100, Vessel Thick Thresh: 1.

**Fig. S9:**
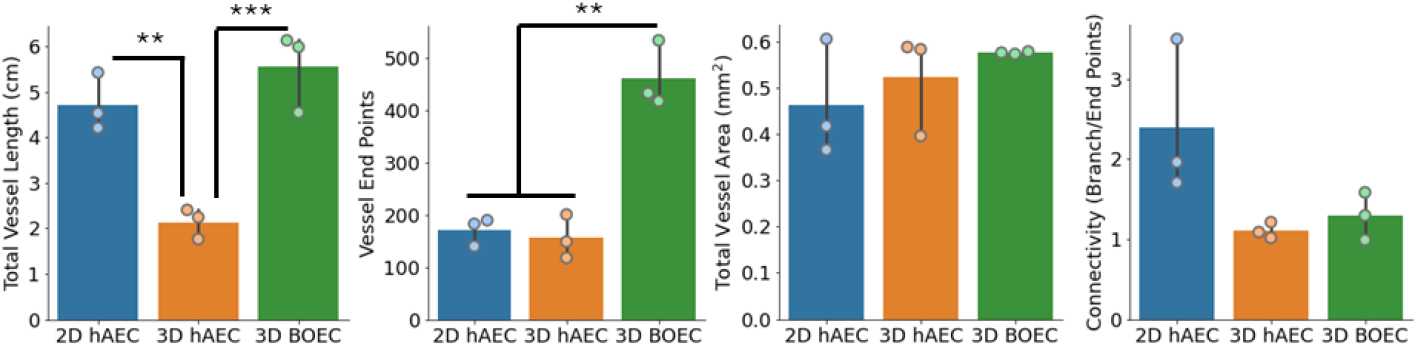
Additional metrics of BRAT characterising diverse microvascular networks. See Fig. 5 for main analysis. *\*p<0*.*05, **p<0*.*01, ***p<0*.*001*.

## References

[1] M.C. Allenby, M.A. Woodruff, Image analyses for engineering advanced tissue biomanufacturing processes, Biomaterials. 284 (2022) 121514. 10.1016/j.biomaterials.2022.121514.

[2] A.R. Murphy, M.C. Allenby, In vitro microvascular engineering approaches and strategies for interstitial tissue integration, Acta Biomater. 171 (2023) 114–130. 10.1016/j.actbio.2023.09.019.

[3] T. Yu, Q. Yang, B. Peng, Z. Gu, D. Zhu, Vascularized organoid-on-a-chip: design, imaging, and analysis, Angiogenesis. 27 (2024) 147–172. 10.1007/s10456-024-09905-z.

[4] Z. Liu, L. Jin, J. Chen, Q. Fang, S. Ablameyko, Z. Yin, Y. Xu, A survey on applications of deep learning in microscopy image analysis, Comput. Biol. Med. 134 (2021) 104523. 10.1016/j.compbiomed.2021.104523.

[5] R. Szeliski, Computer Vision: Algorithms and Applications, 2022.

[6] B. Wang, What’s next for bioimage analysis?, Nat. Methods. 20 (2023) 945–946. 10.1038/s41592-023-01950-8.

[7] A.H. Song, G. Jaume, D.F.K. Williamson, M.Y. Lu, A. Vaidya, T.R. Miller, F. Mahmood, Artificial intelligence for digital and computational pathology, Nat. Rev. Bioeng. 1 (2023) 930–949. 10.1038/s44222-023-00096-8.

[8] A.L. Kiemen, A.M. Braxton, M.P. Grahn, K.S. Han, J.M. Babu, R. Reichel, A.C. Jiang, B. Kim, J. Hsu, F. Amoa, S. Reddy, S.M. Hong, T.C. Cornish, E.D. Thompson, P. Huang, L.D. Wood, R.H. Hruban, D. Wirtz, P.H. Wu, CODA: quantitative 3D reconstruction of large tissues at cellular resolution, Nat. Methods. 19 (2022) 1490–1499. 10.1038/s41592-022-01650-9.

[9] D. and Loncar, C. Mathers, Projections of Global Mortality and Burden of Disease from 2002 to 2030, Plos Med. (2006). https://journals.plos.org/plosmedicine/article?id=10.1371/journal.pmed.0030442.

[10] S. He, G.E. Lim, The Application of High-Throughput Approaches in Identifying Novel Therapeutic Targets and Agents to Treat Diabetes, Adv. Biol. 7 (2023). 10.1002/adbi.202200151.

[11] C.J. Curtaz, S. Wucherpfennig, E. Al-Masnaea, S.-L. Herbert, A. Wöckel, P. Meybohm, M. Burek, High-throughput drug screening to investigate blood-brain barrier permeability in vitro with a focus on breast cancer chemotherapeutic agents, Front. Drug Deliv. 4 (2024) 1–10. 10.3389/fddev.2024.1331126.

[12] S.A. Goel, L.W. Guo, B. Wang, S. Guo, D. Roenneburg, G.E. Ananiev, F.M. Hoffmann, K.C. Kent, High-throughput screening identifies idarubicin as a preferential inhibitor of smooth muscle versus endothelial cell proliferation, PLoS One. 9 (2014) 1–11. 10.1371/journal.pone.0089349.

[13] Niemistö, V. Dunmire, O. Yli-Harja, W. Zhang, I. Shmulevich, Robust quantification of in vitro angiogenesis through image analysis, IEEE Trans. Med. Imaging. 24 (2005) 549–553. 10.1109/TMI.2004.837339.

[14] E. Zudaire, L. Gambardella, C. Kurcz, S. Vermeren, A computational tool for quantitative analysis of vascular networks, PLoS One. 6 (2011) 1–12. 10.1371/journal.pone.0027385.

[15] P. Nowak-Sliwinska, K. Alitalo, E. Allen, A. Anisimov, A.C. Aplin, R. Auerbach, H.G. Augustin, D.O. Bates, J.R. van Beijnum, R.H.F. Bender, G. Bergers, A. Bikfalvi, J. Bischoff, B.C. Böck, P.C. Brooks, F. Bussolino, B. Cakir, P. Carmeliet, D. Castranova, A.M. Cimpean, O. Cleaver, G. Coukos, G.E. Davis, M. De Palma, A. Dimberg, R.P.M. Dings, V. Djonov, A.C. Dudley, N.P. Dufton, S.M. Fendt, N. Ferrara, M. Fruttiger, D. Fukumura, B. Ghesquière, Y. Gong, R.J. Griffin, A.L. Harris, C.C.W. Hughes, N.W. Hultgren, M.L. Iruela-Arispe, M. Irving, R.K. Jain, R. Kalluri, J. Kalucka, R.S. Kerbel, J. Kitajewski, I. Klaassen, H.K. Kleinmann, P. Koolwijk, E. Kuczynski, B.R. Kwak, K. Marien, J.M. Melero-Martin, L.L. Munn, R.F. Nicosia, A. Noel, J. Nurro, A.K. Olsson, T. V. Petrova, K. Pietras, R. Pili, J.W. Pollard, M.J. Post, P.H.A. Quax, G.A. Rabinovich, M. Raica, A.M. Randi, D. Ribatti, C. Ruegg, R.O. Schlingemann, S. Schulte-Merker, L.E.H. Smith, J.W. Song, S.A. Stacker, J. Stalin, A.N. Stratman, M. Van de Velde, V.W.M. van Hinsbergh, P.B. Vermeulen, J. Waltenberger, B.M. Weinstein, H. Xin, B. Yetkin-Arik, S. Yla-Herttuala, M.C. Yoder, A.W. Griffioen, Consensus guidelines for the use and interpretation of angiogenesis assays, Angiogenesis. 21 (2018) 425–532. 10.1007/s10456-018-9613-x.

[16] M.E. Seaman, S.M. Peirce, K. Kelly, Rapid analysis of vessel elements (RAVE): A tool for studying Physiologic, Pathologic and Tumor Angiogenesis, PLoS One. 6 (2011). 10.1371/journal.pone.0020807.

[17] B.A. Corliss, R.W. Doty, C. Mathews, P.A. Yates, T. Zhang, S.M. Peirce, REAVER: A program for improved analysis of high-resolution vascular network images, Microcirculation. 27 (2020) 1–14. 10.1111/micc.12618.

[18] M.B. Vickerman, P.A. Keith, T.L. McKay, D.J. Gedeon, M. Watanabe, M. Montano, G. Karunamuni, P.K. Kaiser, J.E. Sears, Q. Ebrahem, D. Ribita, A.G. Hylton, P. Parsons-Wingerter, VESGEN 2D: Automated, user-interactive software for quantification and mapping of angiogenic and lymphangiogenic trees and networks, Anat. Rec. 292 (2009) 320–332. 10.1002/ar.20862.

[19] J.A. Montoya-Zegarra, E. Russo, P. Runge, M. Jadhav, A.H. Willrodt, S. Stoma, S.F. Nørrelykke, M. Detmar, C. Halin, AutoTube: a novel software for the automated morphometric analysis of vascular networks in tissues, Angiogenesis. 22 (2019) 223–236. 10.1007/s10456-018-9652-3.

[20] X. Wang, G. Zhu, S. Wang, J. Rhen, J. Pang, Z. Zhang, Vessel tech: a high-accuracy pipeline for comprehensive mouse retinal vasculature characterization, Angiogenesis. 24 (2021) 7–11. 10.1007/s10456-020-09752-8.

[21] B. Callewaert, W. Gsell, U. Himmelreich, E.A.V. Jones, Q-VAT: Quantitative Vascular Analysis Tool, Front. Cardiovasc. Med. 10 (2023) 1–11. 10.3389/fcvm.2023.1147462.

[22] Jaeschke, H. Eckert, L.J. Bray, Qiber3D - An open-source software package for the quantitative analysis of networks from 3D image stacks, Gigascience. 11 (2022) 1–9. 10.1093/gigascience/giab091.

[23] S.D. McGarry, C. Adjekukor, S. Ahuja, J. Greysson-Wong, I. Vien, K.D. Rinker, S.J. Childs, Vessel Metrics: A software tool for automated analysis of vascular structure in confocal imaging, Microvasc. Res. 151 (2024) 104610. 10.1016/j.mvr.2023.104610.

[24] Rota, L. Possenti, G.S. Offeddu, M. Senesi, A. Stucchi, I. Venturelli, T. Rancati, P. Zunino, R.D. Kamm, M.L. Costantino, A three-dimensional method for morphological analysis and flow velocity estimation in microvasculature on-a-chip, Bioeng. Transl. Med. 8 (2023) 1–12. 10.1002/btm2.10557.

[25] G.R. Untracht, R.S. Matos, N. Dikaios, M. Bapir, A.K. Durrani, T. Butsabong, P. Campagnolo, D.D. Sampson, C. Heiss, D.M. Sampson, OCTAVA: An open-source toolbox for quantitative analysis of optical coherence tomography angiography images, PLoS One. 16 (2021) 1–22. 10.1371/journal.pone.0261052.

[26] M.C. Allenby, E.S. Liang, J. Harvey, M.A. Woodruff, M. Prior, C.D. Winter, D. Alonso-Caneiro, Detection of clustered anomalies in single-voxel morphometry as a rapid automated method for identifying intracranial aneurysms, Comput. Med. Imaging Graph. 89 (2021). 10.1016/j.compmedimag.2021.101888.

[27] Bhargava, B. Monteagudo, P. Kushwaha, J. Senarathna, Y. Ren, R.C. Riddle, M. Aggarwal, A.P. Pathak, VascuViz: a multimodality and multiscale imaging and visualization pipeline for vascular systems biology, Nat. Methods. 19 (2022) 242–254. 10.1038/s41592-021-01363-5.

[28] B.M. Craver, M.M. Acharya, B.D. Allen, S.N. Benke, N.W. Hultgren, J.E. Baulch, C.L. Limoli, 3D surface analysis of hippocampal microvasculature in the irradiated brain, Environ. Mol. Mutagen. 57 (2016) 341–349. 10.1002/em.22015.

[29] A.R. Murphy, R.A. Franco, M.C. Allenby, Fabricating microfluidic co-cultures of immortalised cell lines uncovers robust design principles for the simultaneous formation of patterned, vascularised, and stem cell-derived adipose tissue., BioRxiv. (2025). 10.1101/2025.01.22.634386.

[30] Z. Jun, H. Jinglu, Image segmentation based on 2D Otsu method with histogram analysis, in: Proc. - Int. Conf. Comput. Sci. Softw. Eng. CSSE 2008, 2008: pp. 105–108. 10.1109/CSSE.2008.206.

[31] S. Van Der Walt, J.L. Schönberger, J. Nunez-Iglesias, F. Boulogne, J.D. Warner, N. Yager, E. Gouillart, T. Yu, Scikit-image: Image processing in python, PeerJ. 2014 (2014). 10.7717/peerj.453.

[32] D. Gupta, R.S. Anand, A hybrid edge-based segmentation approach for ultrasound medical images, Biomed. Signal Process. Control. 31 (2017) 116–126. 10.1016/j.bspc.2016.06.012.

[33] J. Zhang, Z. Lu, M. Li, Active Contour-Based Method for Finger-Vein Image Segmentation, IEEE Trans. Instrum. Meas. 69 (2020) 8656–8665. 10.1109/TIM.2020.2995485.

[34] R.M. Abarca, Network Science with Python and NetworkX Quick Start Guide, 2021.

[35] P.G. Camici, G. D’Amati, O. Rimoldi, Coronary microvascular dysfunction: Mechanisms and functional assessment, Nat. Rev. Cardiol. 12 (2015) 48–62. 10.1038/nrcardio.2014.160.

[36] N. Bhandari, J.D. Newman, J.S. Berger, N.R. Smilowitz, Diabetes mellitus and outcomes of lower extremity revascularization for peripheral artery disease, Eur. Hear. J. - Qual. Care Clin. Outcomes. 8 (2022) 298–306. 10.1093/ehjqcco/qcaa095.

[37] D. Pellegrini, R. Kawakami, G. Guagliumi, A. Sakamoto, K. Kawai, A. Gianatti, A. Nasr, R. Kutys, L. Guo, A. Cornelissen, L. Faggi, M. Mori, Y. Sato, I. Pescetelli, M. Brivio, M. Romero, R. Virmani, A. V. Finn, Microthrombi as a Major Cause of Cardiac Injury in COVID-19 A Pathologic Study, Circulation. 143 (2021) 1031–1042. 10.1161/CIRCULATIONAHA.120.051828.

